# Comparative evaluation of the Geenius™ HIV 1/2 Confirmatory Assay and the HIV-1 and HIV-2 Western blots in the Japanese population

**DOI:** 10.1101/334888

**Authors:** Makiko Kondo, Koji Sudo, Takako Sano, Takuya Kawahata, Ichiro Itoda, Shinya Iwamuro, Yukihiro Yoshimura, Natsuo Tachikawa, Yoko Kojima, Haruyo Mori, Hiroshi Fujiwara, Naoki Hasegawa, Shingo Kato

**Affiliations:** Division of Microbiology, Kanagawa Prefectural Institute of Public Health, Chigasaki, Kanagawa, Japan; Department of Microbiology and Immunology, Keio University School of Medicine, Shinjuku-ku, Tokyo, Japan; Virology Section, Division of Microbiology, Osaka Institute of Public Health, Osaka, Osaka, Japan; Shirakaba Clinic, Shinjuku-ku, Tokyo, Japan; Atsugi City Hospital, Atsugi, Kanagawa, Japan; Department of Infectious Diseases, Yokohama Municipal Citizen’s Hospital, Yokohama,Kanagawa, Japan; Center for Infectious Diseases and Infection Control, Keio University Hospital, Shinjuku-ku,Tokyo, Japan

**Author notes:** Corresponding author (SK).

## Abstract

Accurate diagnosis of earlier HIV infection is essential for treatment and prevention. Currently, confirmation tests of HIV infection in Japan are performed using Western blot (WB), but WB has several limitations including low sensitivity and cross-reactivity between HIV-1 and HIV-2 antibodies. To address these problems, a new HIV testing algorithm and a more reliable confirmation and HIV-1/2 differentiation assay are required. The Bio-Rad Geenius™ HIV-1/2 Confirmatory Assay (Geenius) has recently been approved and recommended for use in the revised guidelines for diagnosis of HIV infection by the Center for Disease Control and Prevention (USA). We made comprehensive comparison of the performance of Geenius and the Bio-Rad NEW LAV BLOT 1 and 2 (NLB 1 and 2) which are WB kits for HIV-1 and HIV-2, respectively, to examine if Geenius is a suitable alternative to these WB assays which are now being used in HIV testing in Japan. A total of 166 HIV-1 positive samples (146 from patients with established HIV-1 infection and 20 from patients with acute infection), five HIV-1 seroconversion panels containing 21 samples and 30 HIV-2 positive samples were used. In addition, a total of 140 HIV negative samples containing 10 false-positives on screening tests were examined. The sensitivity of Geenius and NLB 1 for HIV-1 positive samples was 99.3% and 98.6%, respectively. Geenius provided more positive results in the samples from acute infections and detected positivity 0 to 32 days earlier in seroconversion panels than NLB 1. NLB 2 gave positive results in 12.3% of HIV-1 positive samples. The sensitivity of both Geenius and NLB 2 for HIV-2 positive samples was 100%. The specificity of Geenius, NLB 1 and NLB 2 was 98.5%, 81.5% and 90.0%, respectively. Geenius is an attractive alternative to WB for confirmation and differentiation of HIV-1 and HIV-2 infections. The adaptation of Geenius to the HIV testing algorithm may be advantageous for rapid diagnosis and the reduction of testing costs.

## Introduction

The risk of HIV transmission during acute and early infection is much higher than that during established infection [1]. Furthermore, early initiation of antiretroviral therapy (ART) substantially reduces the risk of transmission to sexual partners [2] and improves clinical outcomes, compared with delayed ART [3]. Accurate diagnosis of earlier HIV infection is important for treatment and prevention strategies.

Currently, diagnosis of HIV infection in Japan is carried out mainly in two different algorithms: (i) a sample tested positive on HIV-1/2 antigen/antibody assay is retested with HIV-1 Western blot (WB-1) and HIV-2 Western blot (WB-2) simultaneously, and then, if the results on both assays are negative, applied to nucleic acid test (NAT) of HIV-1 plasma RNA; this algorithm is recommended by the National Institute of Infectious Diseases (Japan) [4]; (ii) a sample that tested positive on HIV-1/2 antigen/antibody assay is then retested with WB-1 and NAT at the same time, and then, if the results on both assays are negative, applied to WB-2; this is recommended by the Japanese Society for AIDS Research [5]. These algorithms, however, have several limitations associated with Western blot that include false negative or indeterminate results in the early phase, cross-reactivity between HIV-1 and HIV-2 [6], and a labor-intensive and time-consuming protocol.

In 2014, the Center for Disease Control and Prevention (CDC) in the US published revised guidelines for diagnosis of HIV infection in which the use of an HIV-1 and HIV-2 antibody differentiation assay is recommended after a repeatedly reactive HIV-1/2 antigen/antibody test [7]. The FDA-approved Multispot™ HIV-1/HIV-2 Rapid Test (Bio-Rad Laboratories) was initially validated for this purpose. Thereafter, Bio-Rad developed a new confirmatory and differentiation test, the Geenius™ HIV-1/2 Confirmatory Assay (hereafter called Geenius). Geenius received a CE mark in February 2013 and clearance from the Food and Drug Administration in October 2014. Although Geenius has been evaluated in many studies [8–17], there have been few studies on comparison between Geenius and WB. Moon et al. compared the performance of Geenius and WB-1 [16] but did not tested WB-2, and thus they did not comparatively evaluate the HIV-1/2 differentiation ability of Geenius and WB-1/WB-2.

In Japan, while Geenius has not been approved yet, there is a growing interest in the CDC-recommended HIV diagnostic algorithm because it is expected to decrease the number of indeterminate results, allow earlier identification of HIV infections, and reduce the number of NAT to resolve the ambiguity of WB results.

The aims of this study are to compare the confirmation and differentiation performance of Geenius and NEW LAV BLOT 1 and 2 (Bio-Rad Laboratories, Tokyo, Japan, hereafter called NLB 1 and 2), which are WB-1 and WB-2 kits, respectively, and to examine if Geenius is a suitable alternative to WB in the HIV testing algorithm in Japan.

## Material and methods

### Samples and patients

A total of 166 HIV-1 positive samples were used: 146 were obtained from patients with established HIV-1 infection and 20 from patients with acute infection. Among the patients with established infection, 73 were obtained from patients receiving ART at the Keio University Hospital or Atsugi City Hospital and had been diagnosed with HIV-1 infection by either of Dainascreen® HIV Combo (an HIV-1 p24 Ag/HIV-1/2 Ab immunochromatographic test, Alere Medical, Tokyo, Japan) or the Architect® HIV Ag/Ab Combo Assay (an automated HIV-1 p24 Ag/HIV-1/2 Ab test, Abbott Japan, Chiba, Tokyo), followed by NLB 1 and 2 and, if necessary, the Cobas AmpliPrep/Cobas TaqMan® HIV-1 Test (an automated qualitative HIV-1 RNA test, Roche Diagnostics, Tokyo, Japan, hereafter called Cobas). The other 93 samples were obtained from individuals seeking HIV testing in public health centers located in Kanagawa and Osaka: 85 were positive on Dainascreen HIV Combo and 8 were positive on the Architect® HIV Ag/Ab Combo Assay. Their infections were confirmed by NLB 1 and 2 or Cobas. Established HIV-1 infection was defined by positive results on both NLB 1 and Cobas; acute HIV-1 infection was defined by an indeterminate or negative result on NLB 1 but a positive Cobas result.

Five HIV-1 seroconversion panels comprised of seven, five, four, three and two samples, respectively, were obtained from patients attending the Shirakaba Clinic in Tokyo, Japan.

Thirty samples of two commercially obtained HIV-2 panels were used: five from HIV-2 Mixed Titer AccuSet™ Performance Panel (PRE301B, SeraCare Life Sciences, Millford, MA) and 25 from Plasma-CPD-A Anti HIV-2 (HemaCare, Los Angeles, CA).

A total of 140 HIV negative samples were obtained from individuals seeking HIV testing in the public health centers, which were tested as mentioned above; 10 of them were false-positive on screening tests using Dainascreen HIV Combo, which were negative or indeterminate on NLB 1 and 2, and negative on Cobas.

Comparative testing by Geenius and NLB 1 and 2 was conducted between May 2016 and April 2017 in Kanagawa Prefectural Institute of Public Health, Osaka Institute of Public Health, and Keio University School of Medicine according to the manufacturer’s instructions.

### Geenius

Geenius is a single-use immunochromatographic test for the confirmation and differentiation of individual antibodies to HIV-1 and HIV-2 in whole blood, serum or plasma samples using HIV synthetic peptides or recombinant proteins for HIV-1 (p31 [POL], gp160 [ENV], p24 [GAG] and gp41 [ENV]) and HIV-2 (gp36 [ENV] and gp140[ENV]). Geenius is aimed at confirming the presence of antibodies to HIV-1 and HIV-2 in samples reactive by screening tests. Banding patterns and intensities on a Geenius cassette were read by an automated reader connected to a personal computer and interpreted using the Geenius algorithm. This cartridge assay allows rapid evaluation within 30 min. Interpretive results involve HIV negative, HIV-2 indeterminate, HIV-1 indeterminate, HIV indeterminate, HIV-1 positive, HIV-2 positive, HIV-2 positive (with HIV-1 cross-reactivity), and HIV positive untypable.

### NLB 1 and 2

NLB 1 and 2 are the only WB kits approved by The Pharmaceuticals and Medical Devices Agency (PMDA) of Japan for confirmation of HIV-1 and HIV-2 infection, respectively. Bands were observed visually. Interpretation of banding patterns is performed as follows: for HIV-1, the presence of at least two of three ENV bands (GP160, GP120 and GP41) is considered positive, no HIV-1 specific band negative, and other patterns indeterminate; for HIV-2, the presence of one ENV, one GAG and one POL band is considered positive, no HIV-2 specific band negative, other patterns indeterminate.

### Statistics

Sensitivity and specificity were determined by considering indeterminate results as not positive and not negative, respectively, with 95% confidence interval [95% CI]. Cohen’s kappa (κ) was calculated to assess agreement between Geenius and NLB 1.

## Results

### Samples in established HIV-1 infection

Geenius, NLB 1, and NLB 2 results on 146 samples from patients with established HIV-1 infection are compared in Table 1. Geenius provided 145 HIV-1 positive results including one HIV positive untypable (sensitivity, 99.3% [95% CI, 96.2–99.8]) and one HIV-1 indeterminate result. NLB 1 showed 144 positive result (sensitivity, 98.6% [95% CI, 95.1–99.6]) and two indeterminate results: the indeterminate results were observed on samples from patients receiving ART. It is notable that only four samples were negative by NLB 2, which may be due to high cross-reactivity.

**Table 1.**
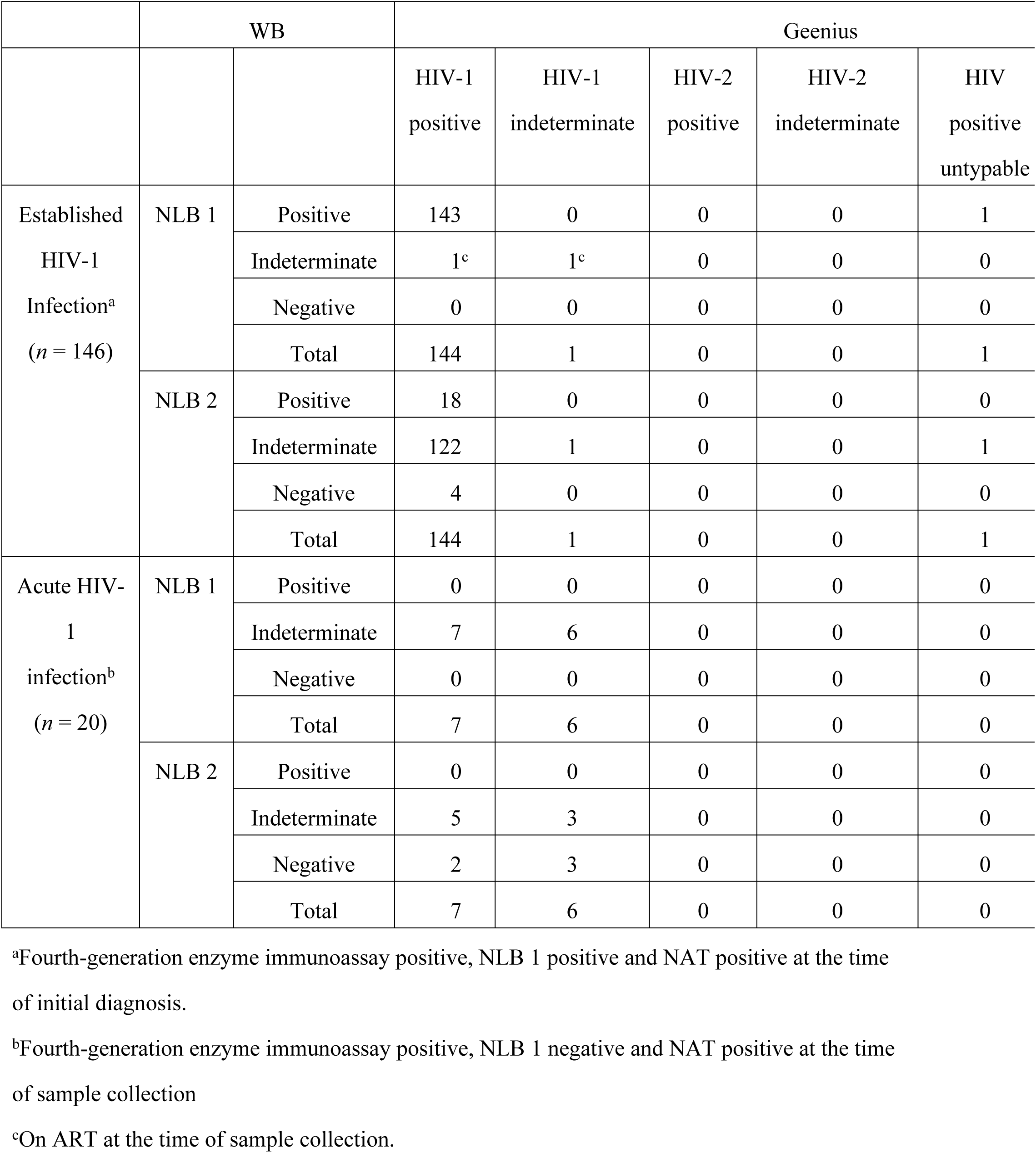
Comparison of Geenius with NLB 1 and 2 results for established and acute HIV-1 infection samples.

### Samples in acute HIV-1 infection

Geenius, NLB 1, and NLB 2 results on 20 samples from patients with acute HIV-1 infection are compared in Table 1. Geenius reclassified seven of the NLB 1 indeterminate samples as positive, showing that Geenius has a higher detection sensitivity than NLB 1.

### Seroconversion panels

Five HIV-1 seroconversion panels were used to compare the detection ability of identifying positive samples during the early phase of infection between Geenius and NLB 1 (Table 2). Geenius gave positive results 0 to 32 days earlier than NLB 1. Cross-reactive p26 bands appeared in NLB 2 as the specific HIV-1 antibody titer increased, while no HIV-2-related band was observed in Geenius.

**Table 2.**
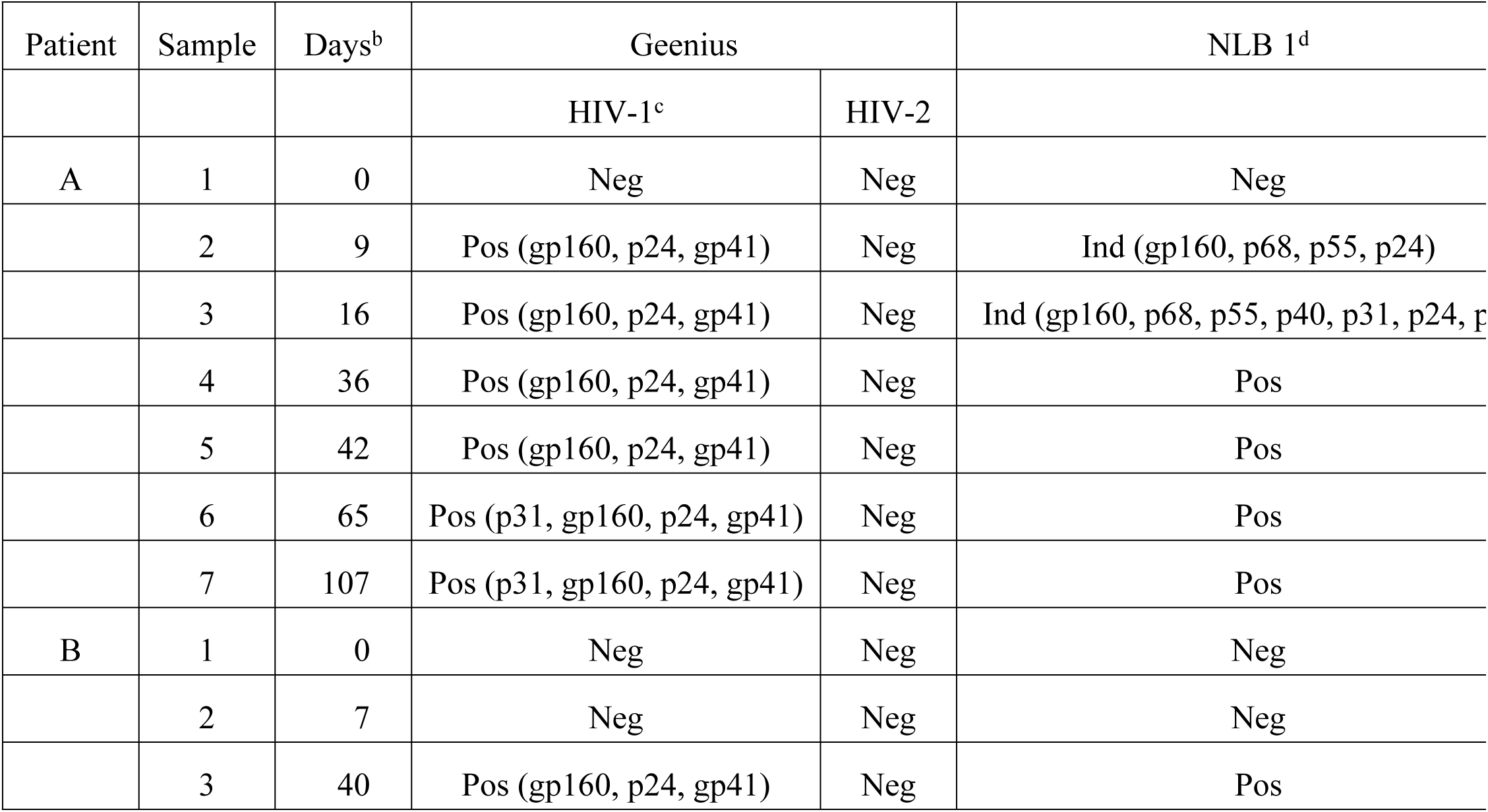
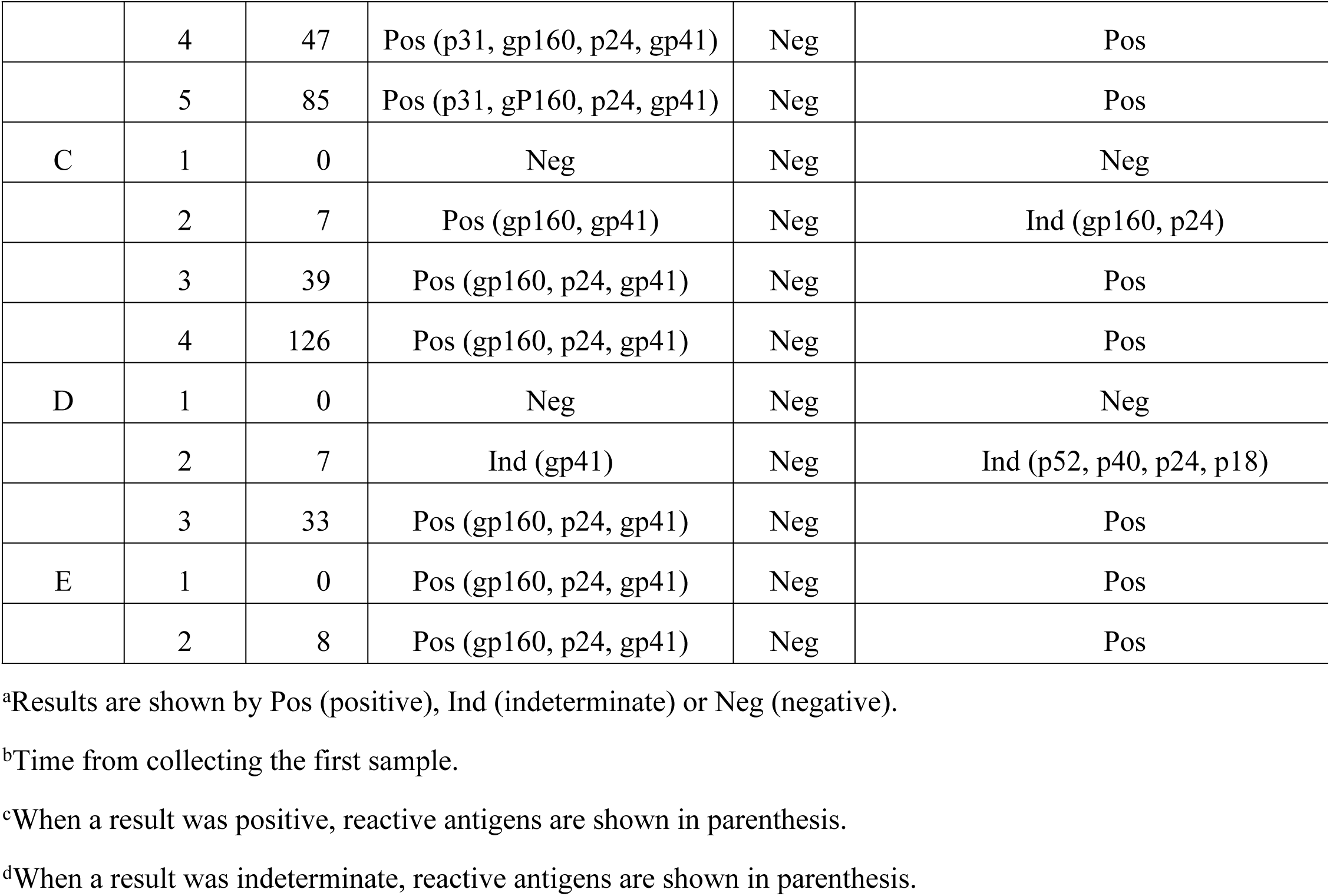
Comparison of Geenius with NLB 1 and 2 results for HIV-1 seroconversion panels^a^.

### HIV-2 panels

Thirty samples of two commercial HIV-2 panels were used to compare Geenius, NLB 1, and NLB 2 (Table 3). All samples were positive with NLB 2 (sensitivity, 100% [95% CI, 88.4– 99.5]); two samples were positive and 28 were indeterminate with NLB 1 (false-positive rate, 6.7% [95% CI, 2.1–12.1]). Geenius gave 28 HIV-2 positive and two HIV positive untypable results (sensitivity, 100% [95% CI, 88.4–99.5]).

**Table 3.** Comparison of Geenius with NLB 1 and 2 results for HIV-2 panel samples.

|  |  | Geenius |  |  |  |
| --- | --- | --- | --- | --- | --- |
|  |  | HIV-1 | HIV-2 | HIV-2 positive with HIV-1 | HIV positive |

|  |  | positive | positive | cross-reactivity | untypable | ne |
| --- | --- | --- | --- | --- | --- | --- |
| NLB 1 | Positive | 0 | 0 | 2 | 0 |  |
|  | Indeterminate | 0 | 10 | 16 | 2 |  |
|  | Negative | 0 | 0 | 0 | 0 |  |
|  | Total | 0 | 10 | 18 | 2 |  |
| NLB 2 | Positive | 0 | 10 | 18 | 2 |  |
|  | Indeterminate | 0 | 0 | 0 | 0 |  |
|  | Negative | 0 | 0 | 0 | 0 |  |
|  | Total | 0 | 10 | 18 | 2 |  |

### Seronegative samples

A total of 130 screening negative samples were used to determine the specificity of three assays (Table 4). Concordant negative results between Geenius and NLB 1 were obtained for 104 samples; those between Geenius and NLB 2 for 116 samples. The specificity of Geenius, NLB 1, and NLB 2 were 98.5% (128/130) [95% CI, 94.6–99.5], 81.5% (106/130) [95% CI, 73.8–87.2] and 90.0% (117/130) [95% CI, 83.5–94.0], respectively.

**Table 4.** Comparison of Geenius with NLB 1 and 2 results for negative samples by fourth-generation immunoassay (*n* = 130).

|  |  | Geenius |  |  |  |  |
| --- | --- | --- | --- | --- | --- | --- |
|  |  | HIV-1<br>positive | HIV-1<br>indeterminate | HIV-2<br>positive | HIV-2<br>indeterminate | HIV positive<br>untypable |
| NLB 1 | Positive | 0 | 0 | 0 | 0 | 0 |
|  | Indeterminate | 0 | 0 | 0 | 0 | 0 |
|  | Negative | 0 | 1 | 0 | 1 | 0 |
|  | Total | 0 | 1 | 0 | 1 | 0 |
| NLB 2 | Positive | 0 | 0 | 0 | 0 | 0 |
|  | Indeterminate | 0 | 0 | 0 | 1 | 0 |
|  | Negative | 0 | 1 | 0 | 0 | 0 |
|  | Total | 0 | 1 | 0 | 1 | 0 |

### False-positive samples

It is important for a confirmatory assay to discriminate between acute HIV-1 infections and false positive screening results. Ten Dainascreen HIV Combo positive but Cobas negative samples were tested with the three assays (Table 5): eight were negative and two were indeterminate (positive p31 bands) with Geenius; six were negative and four were indeterminate with NLB 1; five were negative and five were indeterminate with NLB 2, suggesting Geenius is the most specific for HIV-1 false-positive screening samples among the three kits.

**Table 5.** Comparison of Geenius with NLB 1 and 2 results for HIV-1 Combo positive but NAT negative samples (*n* = 10)^a^.

| Sample. | Geenius<sup>a</sup> |  | NLB 1 |  |
| --- | --- | --- | --- | --- |
|  | HIV-1 | HIV-2 |  |  |
| 1 | Negative | Negative | Indeterminate (p18) |  |
| 2 | Negative | Negative | Indeterminate (p18) | Indeter |
| 3 | Negative | Negative | Negative |  |
| 4 | Negative | Negative | Negative |  |
| 5 | Negative | Negative | Indeterminate (p24, p18) |  |
| 6 | Indeterminate (p31) | Negative | Indeterminate (p18) | Inde |
| 7 | Negative | Negative | Negative |  |
| 8 | Negative | Negative | Negative | Inde |
| 9 | Indeterminate (p31) | Negative | Negative | Inde |
| 10 | Negative | Negative | Negative | Inde |

### Concordance

The overall concordance (*κ*) between Geenius and NLB 1 was 0.78 if positive, indeterminate, and negative results were considered separately, and 0.95 if indeterminate results were considered as negative.

## Discussion

Japan is a country with low-level HIV epidemics. The cumulative reported incidence of HIV infection through the end of 2016 was 27,443 [18]. Among them, the number of persons with HIV-2 infection was six [19–22], and there has been no report of HIV-1 and HIV-2 dual infection. According to PMDA, the confirmation of HIV-1 and HIV-2 infections should be performed using WB-1 and WB-2, respectively. However, discrimination between HIV-1 and HIV-2 infections is sometimes very difficult due to cross-reactivity of antibodies against the two viruses. In such cases, it is recommended that the samples are retested from a screening test after several weeks or tested with SERODIA®-HIV-1/2 (a particle agglutination assay to detect antibodies to HIV-1 and/or HIV-2, Fujirebio, Tokyo, Japan) or Pepti-LAV 1/2 Assay (an enzyme immunoassay for differentiation of HIV-1 and HIV-2 antibodies, Bio-Rad, Tokyo, Japan) to distinguish HIV-1 and HIV-2 infections, while these differentiation assays also have a high cross-reactivity. These additional tests are, however, laborious, time-consuming, and costly, and cause a large burden in countries such as Japan where the prevalence of HIV-2 infection is extremely low. In this study, we aimed to assess whether a new rapid test Geenius is an effective alternative to WB-1 and WB-2 for confirmation and discrimination of HIV-1 and HIV-2 infections.

Although the sensitivity of Geenius and NLB 1 was not significantly different (99.3% vs 98.6%) for samples from established HIV-1 infections, Geenius gave seven positive results in 20 NLB 1 negative or indeterminate samples from acute HIV-1 infections and provided positive results earlier than NLB 1 in two of five seroconversion panels, showing that Geenius is more sensitive than NLB 1. For 140 HIV-1 negative samples including 10 false-positive samples, Geenius gave 136 negative and NLB 1 gave 112 negative results, showing that Geenius is more specific than NLB 1.

Cross-reactivity of HIV-1 and HIV-2 antibodies between NLB 1 and NLB 2 was remarkable compared with Geenius. When HIV-1 positive samples were examined, 18 of 144 NLB 1 positive samples were also positive with NLB 2. Geenius, however, resolved all of these double-positive samples as HIV-1 positive. An overall discrimination rate of Geenius was 97.7% (172/176) [95% CI, 94.3–99.1] and that of a combinational use of NLB 1 and NLB 2 was 87.5% (154/176) [95% CI, 81.7–91.5], showing that Geenius has a higher discrimination ability than NLB 1/NLB 2. Geenius still gave three HIV positive untypable results: one in 146 HIV-1 positive samples and two in 20 HIV-2 positive samples. It is practically impossible to determine if these results reflect HIV-1/2 dual infection or cross-reactivity at present because the application of HIV-2 NAT for confirmation of HIV-2 infection has not yet been established.

According to the HIV diagnostic algorithm recommended by CDC, samples that are positive on screening tests but negative or indeterminate on HIV-1/HIV-2 antibody differentiation immunoassay should be tested with an HIV-1 NAT [7]. Because Geenius gave fewer negative or indeterminate results than NLB 1/NLB 2 in HIV-1 positive and HIV-1 false-positive samples (Tables 1, 2, and 5), the use of Geenius will decrease the number of required HIV-1 NAT compared to NLB 1/NLB 2, which may lead to the reduction of testing costs.

Geenius is characterized by the cassette involving immunochromatographic components to detect HIV-1/2 antibodies and the automated reader using the proprietary interpretive software. These devices make Geenius have several advantages over WB, including a simple, easy and rapid procedure (within 30 min) and objective interpretation of banding patterns. It is well known that technical skills and interpretation experience are required to perform WB. The rapidity of Geenius may allow HIV testing in public health centers or outreach services to be completed on the same day.

WB is frequently used for estimating the stage of early HIV-1 infections [23], based on the study by Fiebid et al. [24], in which positive WB without p31 band is stage V and positive WB with p31 band is stage VI. In this study, Geenius was shown to confirm HIV-1 seropositivity earlier than WB, and thereafter detect p31 bands in panels A and B (Table 2). Keating et al. [25] demonstrated that additional interpretive analysis of band intensities help estimation of recent infections. Development of such algorithms may contribute to epidemiological studies on HIV infections.

## Conclusions

Geenius is an attractive alternative to WB for confirmation and differentiation of HIV-1 and HIV-2 infections. The adaptation of Geenius to the HIV testing algorithm may lead to a more rapid diagnosis and cost reduction.

## Acknowledgements

The authors would like to thank Bio-Rad Laboratories (Tokyo, Japan) for the supply of free assay kits.

